# What drives wolf preference towards wild ungulates? Insights from a multi-prey system in the Slovak Carpathians

**DOI:** 10.1101/2022.03.02.482693

**Authors:** Nuno F. Guimarães, Francisco Álvares, Jana Ďurová, Peter Urban, Jozef Bučko, Tomáš Iľko, Jaro Brndiar, Jozef Štofik, Tibor Pataky, Miroslava Barančeková, Rudolf Kropil, Peter Smolko

## Abstract

The wolf is a generalist-opportunistic predator that displays diverse and remarkably adaptable feeding strategies across its range with local adaptations to certain prey species depending on their availability and vulnerability. The multi-prey system of the Slovak Carpathians supports important portion of the European wolf population; however, it has been markedly understudied. We evaluated winter diet composition and prey selection of Slovak wolves based on 321 scat samples collected between September – February within four different study areas during 2015 – 2017. The winter diet of wolves in the Slovak Carpathians was characterized by a 98% occurrence of wild large-sized and medium-sized ungulates with red deer occurring in wolf scats most often, consistent with their highest density among other wild ungulates. However, by comparing the consumption with availability of wild prey, we found that wolves in fact selected for wild boar especially in areas with higher altitudinal range, while selected for red deer in areas with low altitudinal range where this prey species was more spatially predictable. Although wolves showed the potential to switch between red deer and wild boar when their density increases, we found that this variation can be rather linked to changing prey vulnerability, which is dependent on particular environmental conditions at local scale such as topography and snow accumulation. The present study provides valuable insights into the winter foraging ecology of Slovak wolves in a multi-prey system of the Carpathians and allows for practical implications in the management of the rapidly increasing populations of wild ungulates across Europe.

## Introduction

The wolf (*Canis lupus*) is the most widespread large carnivore in the world and one of the most important apex-predators with stabilising and sanitary role in the ecosystem [1–3]. Being a generalist-opportunistic predator [4], as a result of its adaptability to different environments, wolf displays diverse and adaptable feeding strategies across its range [5,6] that follow a geographic pattern [7,8]. Wolf prey mainly on large-sized wild ungulates such as moose (*Alces alces*) and reindeer (*Rangifer tarandus*) in northern Europe [7,9] and red deer (*Cervus elaphus*) in central and eastern Europe [10]. In contrast, medium-sized ungulates such as wild boar (*Sus scrofa*) and roe deer (*Capreolus capreolus*) are more typical prey in southern Europe [11–13]. In areas with low availability of wild ungulates, wolf may consume small-sized wild mammals, fish and birds [5,8] but they also turn to anthropogenic resources, such as livestock [12,14,15]. In this context, wolf damage to livestock production, as well as wolf depredation on hunting game species, is a constant source of conflict regarding coexistence with human activities [16–18].

Despite the general geographic pattern in wolf diet throughout Europe, wolf may show local adaptations to other prey species depending on the availability of prey [19] [20] and environmental conditions [21,22]. For example, opportunistic predators living in multiple prey systems tend to select the most abundant prey (*apostatic selection*) [23], and the pattern of selection is influenced by changes in prey availability [24,25]. As a result, wolves show prey switching behaviour [26] between prey species, which reduces predation rate on a particular species at low density and therefore can have a stabilizing effect on the system [27,28]. Wolves hunt any vulnerable prey available in their territory [29]; however, in multiple prey systems wolves often show a clear selection for a single prey species [19,24,30]. According to the optimal diet theory [31](Stephens and Krebs 1986), wolf select the more profitable prey, where profitability is the ratio between energy gain and handling time. Wolf prey species have effective physical and behavioural defence traits, and each prey species requires different amount of effort to be killed [29]. In this context, prey profitability, and, consequently, prey use and selection, are the result of searching time, encounter rate, capture success and risk of injury [24]. Prey vulnerability, i.e., the physical, behavioural and environmental factors that influence the susceptibility to predation [4,32], strongly affects capture success, and consequently handling time [24,25]. Among the factors determining prey vulnerability, body size is the most important [29]; however, environmental conditions may also affect the effort of the predator to encounter prey, and the efficiency by which the prey animal can escape or attack [25,33,34]. For example, snow had a strong effect on mortality of wild boar in the Bialowieza National Park, Poland [35], and previous studies in Scandinavia have shown that increased snow depth resulted in both a higher proportion of moose calves being killed [36] as well as reduced chase distances of wolves on moose and roe deer [37]. In this context, prey density itself may not be a constant clue for determining prey selection.

Although studies on diet and prey selection of European wolves have been conducted extensively [7,38,39], there are still regions with poor knowledge on wolf trophic ecology, as in the case of the Slovak Carpathians. Slovak wolf population is part of the larger Carpathian population and consists of ∼340-450 wolves [40]. The Carpathian Mountains represent one of the largest wolf refuge areas in Europe and are regarded as being of particular importance for the long-term survival of this species in Europe because of their size and potential to serve as a link between northern and southern populations [40,41]. However, wolves in Slovakia suffered a dramatic persecution until 1975 [16,42,43] when wolf gained a partial protection but was still regularly hunted [44]. With Slovakia joining the EU in 2005, wolves became a protected species under the Habitat Directive 92/43/CEE. However, being included in the annex V of the Habitat Directive, wolves in Slovakia were still hunted within annual harvest quotas ranging between 28 – 149 individuals until the 2020/2021 hunting season [45]. In 2021, the Ministry of the Environment of the Slovak Republic listed the wolf as a fully protected species under the implementation of the Decree no. 170/2021, amending the Nature and Landscape Protection Act no. 543/2002. However, wolf remains a highly controversial species, especially among hunters and stakeholders, due to depredation on game species and damage to livestock [1,46]. As a result, there is the need for updated knowledge on wolf diet composition and prey selection in Slovakia, particularly focusing on wild and domestic ungulates.

In this study, we analysed wolf diet and prey selection during winters 2015 – 2017 in the multiple prey system of the Slovak Carpathians, because studies from this area are relatively scarce [47,48]. In particular, (1) we first evaluated the diet composition of wolves in four different study areas within Slovakia. Then, (2) we evaluated the response of wolves to wild ungulate density variations, by calculating both the true selection (<u>*sensu*</u> Levin’s) [49] and the latent selection [50,51], including environmental factors affecting prey mobility such as snow depth and driving forage availability to ungulates such as elevation [52]. We expected that wild ungulates, particularly red deer, would be the most used prey species [47,48], although, we also expected some variation between study areas. We also assumed that wild boar is more negatively affected by snow accumulation than red deer, because of their smaller body size and a greater hindrance of their mobility [53]. Deep snow makes foraging energetically costly and difficult for wild boar, causing starvation and rapid deterioration, especially at high elevations where forage is scarce and difficult to access [54]. Foraging behaviour of predators drives predator–prey dynamics and its understanding is fundamental not only for a suitable management and conservation of both large carnivores and wild ungulates but also for mitigation of existing conflicts with humans.

## Material and Methods

### Study area

We conducted our study in temperate forests region of the central and eastern Slovakia (Fig 1a). Data on wolf diet composition were collected within four different mountain ranges included in the Carpathians and comprising a total area of 1375 km^2^. The Poľana Protected Landscape Area (hereafter Poľana PLA; N48°40’, E19°28’), Vepor Mountains (hereafter Vepor Mts; N48°38’, E19°44’) and Muráň Plateau National Park (hereafter Muráň Plateau NP; N48°45’, E20°0’) are located within central Slovakia, while the Poloniny National Park (hereafter Poloniny NP; N49°06', E 22°17) is located in eastern Slovakia adjacent to Polish and Ukrainian border (Fig 1a). Forests are the dominant land cover type, which in low altitudes are composed mostly of European beach (*Fagus sylvatica*), with admixture of European hornbeam (*Carpinus betulus*), European ash (*Fraxinus excelsior*) and Sycamore maple (*Acer pseudoplatanus*), while in higher altitudes (> 1000 m a.s.l.) Norway spruce (*Picea abies*) is a dominant species, with admixture of European larch (*Larix decidua*), Scots pine (*Pinus sylvestris*) and Silver fir (*Abies alba*).Main wild prey species, such as red deer, wild boar and roe deer are present in all four study areas. Besides the wolf, brown bear (*Ursus arctos*) and Eurasian lynx (*Lynx lynx*) are also present throughout all study areas. The most common livestock species under extensive grazing are sheep (*Ovis aries*), cattle (*Bos taurus*), and goat (*Capra hircus*).

**Fig 1.**
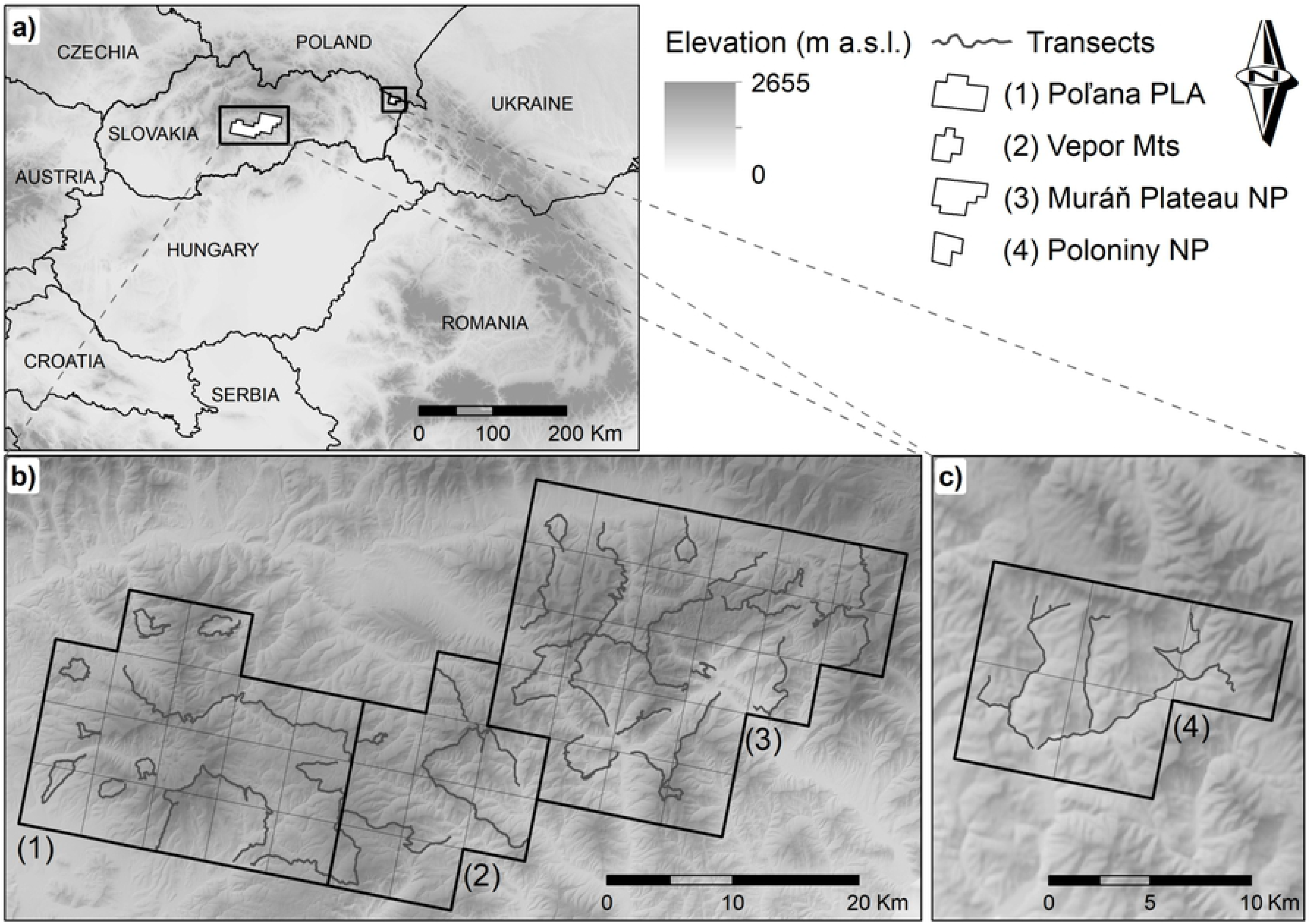
This is the Fig 1 Title: Study area. This is the Fig 1 legend: Figure 1. Location of the four study areas where data for wolf diet analysis were collected within Slovak Carpathians (a), and the network of transects used for winter scat collection between 2015 – 2017 in each study area (b and c). Study areas: Poľana PLA (1), Vepor Mts (2), Muráň Plateau NP (3) and Poloniny NP (4).

### Poľana Protected Landscape Area

The study area in the Poľana PLA, located in the Western Carpathians, encompassed of 425 km^2^ with elevation ranging between 338 – 1458 m a.s.l. (Fig 1b). The region is comprised by 14% deciduous, 24% coniferous and 39% of mixed forests with 3% of shrubs, 9% of pastures, 10% of agricultural land and 1% of human settlements [55]. The annual mean temperature is 6.9 °C (July 18.3 °C and January −3.0 °C), the annual rainfall is 1120 mm, and snow depth > 20 cm covered 50.5% of the area [56]. The Poľana PLA is a UNESCO Biosphere Reserve with restricted agricultural and forestry activities.

### Vepor Mountains

The study area in the Vepor Mts, located in the Western Carpathians, encompassed 225 km2 with elevation ranging between 338 – 1322 m a.s.l. (Fig 1b). The region is comprised by 23% deciduous, 18% coniferous and 33% of mixed forests with 4% of shrub, 14% of pastures, 7% of agricultural land and 1% of human settlements [55]. The annual mean temperature is 7.0 °C (July 18.0°C and January −3.0 °C), the annual rainfall is 1112 mm, and snow depth > 20 cm covered 44.7% of the area [56]. Vepor Mts is exploited from forestry and agricultural perspective with no protection status.

### Muráň Plateau National Park

The study area in the Muráň Plateau NP, located in the Western Carpathians, encompassed 600 km^2^ with elevation ranging between 335 – 1439 m a.s.l. (Fig 1b). The region is comprised by 30% deciduous, 18% coniferous and 28% of mixed forests with 7% of shrub, 7% of pastures, 9% of agricultural land and 1% of human settlements [55]. The annual mean temperature is 6.7 °C (July 17.5°C and January −3.7 °C), the annual rainfall is 965 mm, and snow depth > 20 cm covered 48.0 % of the area [56]. There is a population of ~50 horses that are raised under extensive grazing and trained for work in forestry within the Muráň Plateau NP, although forestry and agriculture are restricted because of the protection status.

### Poloniny National Park

The study area in the Poloniny NP, the only sampling region located in the Eastern Carpathians, encompassed 125 km^2^ with elevation ranging between 338 – 1150 m a.s.l. (Fig 1c). The region is comprised by 80% deciduous, 1% coniferous and 7% of mixed forests with 3% of shrub, 7% of pastures, 2% of agricultural land and no human settlements [55]. The annual mean temperature is 7.0 °C (July 18.0°C and January −3.4°C), the annual rainfall is 725 mm, and snow > 20 cm deep covered 42.2% of the area [56]. As a national park and a valuable source of water, this area is highly protected with minimal agricultural and forestry activities. A population of ~40 European bison (*Bison bonasus*) roam the Poloniny NP, and there is also steady presence of the Eurasian beaver (*Castor fiber*) in the area.

### Scat collection and laboratory analysis

Wolf scats were collected from September to February, between 2015 and 2017. All four sampling areas were divided into 5 × 5 km grid (adapted from 10 km^2^ grid) [57] over a topographical map of the areas. Scat samples were collected within a systematic ground tracking surveys along 487.6 km of randomly selected transects including roads, hiking trails, wildlife trails, and mountain ridges (Fig 1b and 1c). We collected a minimum of 40 wolf scats per sampling area, since according to some studies, this is the minimum systematic sample of scats to be representative of a populations’ diet, rather than a larger sample size that is randomly collected [58,59]. In order to verify the accuracy of the species identification in collected samples, we genetically analysed 53 fresh scats. Species determination was successful in 29 samples, of which 26 (90%) were from wolf while 3 were from red fox (*Vulpes vulpes*) and were removed from further analysis.

Each scat was washed under tap water using a sieve with 1 mm mesh to retain all hairs and other undigested remains (i.e., bones, hooves, feathers) and was then spread on a petri dish and oven dried at 65 C° for 3 to 5 hours [60,61]. Samples lacking hairs or those at high stage of decomposition were discarded (*n* = 6). Blind tests were applied on randomly selected samples from the available collection of hairs of wild and domestic mammal species present in our study area to train and assess the ability of the two observers to identify prey species [6,62], and a species was considered to be accurately determined if the responses of both observers matched in 95% of cases. Identification of each prey species from hair was done systematically using the point frame method [59,60,63] and was based on morphologic features of hairs, such as cuticular pattern, medulla and cross-section [61,64], using available hair identification keys [64–68]. Microscopic analysis was done using a portable microscope MEOPTA model BC 28 SV and Leica dm4000 + camera Leica dfc290 HD.

### Prey availability

Annual data on the availability of wild ungulates were obtained from approved management plans that recorded annual counts of red deer, roe deer and wild boar within mandatory linear surveys conducted by hunters during spring (April). We used the estimates of wild prey (males + females + offspring) between 2015 and 2017 from 22 hunting grounds (20 – 75 km2) within our study areas to spatially map the average annual red deer, wild boar and roe deer population density (Fig 2a; S1 Table). Data on the abundance of wild ungulates in Slovakia was probably not as reliable as scientific surveys of wild ungulates, but since relative differences were important for this analysis, like in Smolko et al. [69,70], we were confident in rough estimates of wild ungulate abundance. Densities of wild ungulates in our study areas ranged between 1.3 – 1.9 ind./km^2^ for red deer, 1.1 – 2.0 ind./km^2^ for roe deer and 0.4 – 0.7 ind./km^2^ for wild boar (Fig 2a).

**Fig 2.**
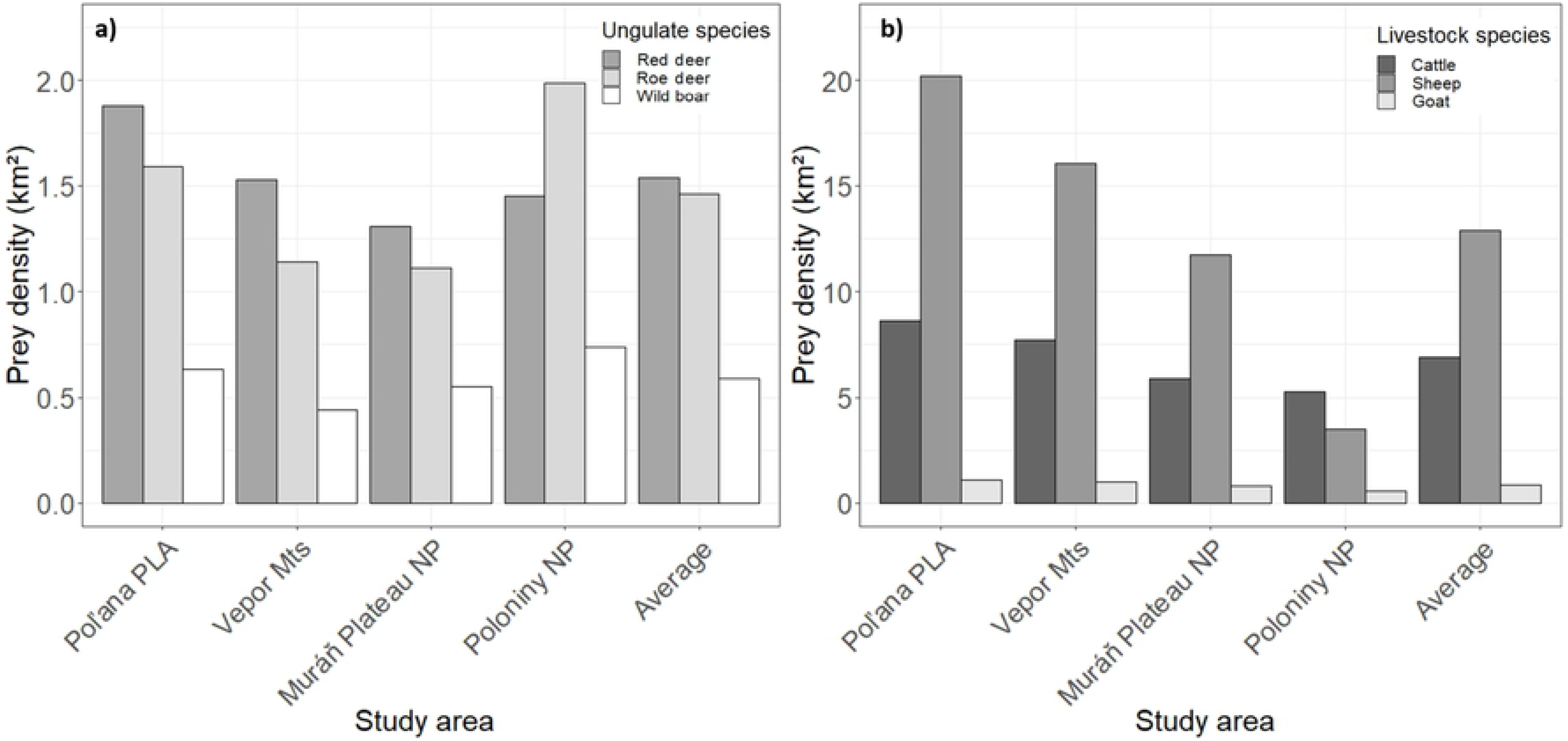
This is the Fig 2 Wild ungulates and livestock densities. This is the Fig 2 legend: Figure 2. Average densities (2015 – 2017) of wild ungulates (a) and livestock (b) in each of the four study areas in the Slovak Carpathians.

Livestock are traditionally raised under extensive grazing on mountain meadows and pastures from April to November [48], being usually under the constant supervision of shepherds with guarding dogs during the day and kept inside enclosures at night. In December however, livestock is brought to low elevations and winters in barns until spring, thus usually not being available as a wolf prey despite having much higher densities compared to wild ungulates (Fig 2b). Annual data on livestock for 2015 – 2017 were obtained from the Statistical Office of the Slovak Republic. Since data on livestock were reported only per municipalities (447 – 1470 km^2^), we calculated average densities for all 9 municipalities within our study areas and created maps for each livestock species (S2 Table). To estimate the wild prey and livestock densities in our study areas we used Zonal statistics tool in ArcMap 10.5 [76] and calculated spatially weighted means per each area and species. We converted prey densities into biomass available per each prey following Ruehe et al [39].

### Data analyses

We calculated the percentage of frequency of occurrence (hereafter *FO*) for each prey item in wolf diet based on the equation: %*FO* = (*N*_*i*_ ÷ *N*_*t*_) × 100, where *N*_*i*_ is the number of occurrences of food item “*i*” and *N*_*t*_ the total number of occurrences of all food items [60].

Considering that degree of digestibility is different for every food item [60], and that *FO* overestimates the importance of prey when the proportion of samples containing only one item is high, we used correction factor (*Y*) developed by Ruehe et al [39] for European ungulates [38] to determine the percentage of ingested biomass (hereafter *BM*). We calculated *BM* following this sequence of three equations: (1) *Y* = 0.00554 + 0.00457 × *X*, where *Y* represents the fresh mass (kg) of prey per scat, and *X* is the average mass of live prey (S2 Table), (2) *BM*_*i*_ = Y × N where *Y* is the digestible biomass of individual type of prey and *N* is the number of samples where we identified the prey, and finally (3) *BM* = *BM*_*i*_ ÷ (*Total BM*_*i*_ × *BM*_*t*_ × 100), where *BM*_*i*_ represents the ingested biomass of species *i*, and *BM*_*t*_ represents the ingested biomass of all species. We used χ^2^-test [71] in order to compare differences in *FO* and *BM* using R software [72]. To evaluate the importance of each prey as a food source, we categorized all prey species into four groups according to Ruprecht [73]: basic (*FO* > 20%); constant (5% < *FO* < 20%); supplementary (1% < *FO* < 5%); and opportunistic (*FO* < 1%). Dietary diversity between study areas was assessed based on the standardized Levin’s formula for measuring the niche breadth [74] using the equation: BS = [(Σp_i_^2^)^−1^ - 1] ÷ (N – 1), where *N* is the number of prey species identified and *p_i_* the proportion of each prey item. A value of, or close to 0 represents a narrow niche breadth (or a high degree of specialization), while a value close, or equal to 1 represents a broad niche breadth (or that the species is a generalist).

To verify if wolves exhibited selection or avoidance of any of the wild ungulate and livestock species present in the habitat, we calculated Ivlev’s electivity index modified by Jacobs [75]: , *D* = (*r*_*i*_ – *p*_*i*_) ÷ [(*r*_*i*_ × *p*_*i*_) - 2*r*_*i*_*p*_*i*_)], where *r*_*i*_ is the relative proportion of *BM* of food item “*i*” and *p_i_* the relative proportion of biomass available of the food item “*i*” in the study area. The values of the index range between −1 and 1, with negative values indicating prey avoidance or inaccessibility, zero showing that prey is randomly consumed, and positive values indicating wolves are actively selecting a specific prey.

In order to test our hypothesis that landscape topography might also affect wolf diet instead of exclusively prey availability, we used a logistic regression to estimate the coefficients of a latent selection difference (LSD) function [50,51]. By using the LSD function, we contrasted prey use of the three most abundant wild ungulates in Slovakia i.e., red deer, wild boar and roe deer. First, we compared red deer (1) with wild boar and roe deer (0), then wild boar (1) with red deer and roe deer (0), and finally roe deer (1) with red deer and wild boar (0) (S3 Table). We tested whether wolves select one prey species over others based on density of prey (ind./km^2^), elevation, study area and proportion of the area within 1 km radius around scat location that was covered with snow >20 cm deep, because locomotion of wild ungulates as well as access to forage are known to become restricted at this depth [30,54]. Elevation model was obtained from the European Environment Agency in 100 × 100 m resolution. Snow depth was obtained from the Carpatclim [56] in 10 x 10 km resolution and interpolated to 100 × 100 m using kriging in the ArcMap software (version 10.5.1) [76]. Next, we used the Focal statistics tool of ArcMap [76] to calculate the proportion of the area covered by snow deeper than 20 cm. We averaged all variables within 1 km-radius (~3 km^2^) representing the smallest size of the core area within wolf home range reported in Europe [77]. Collinearity of the predictor variables was tested by the Pearson’s correlation, while the Akaike’s Information Criterion (AIC_c_) for small sample sizes [78] was used to select the models best explaining wolf prey selection. We assessed the ability of the models to accurately predict wolf winter diet by the k-fold cross-validation to evaluate how well selection models predicted the use of prey by wolf while Spearman’s rank correlation was used to assess the relationship between predicted values and observed frequencies of locations within 10 bins sites with equal areas [79].

## Results

In total, we collected 330 scats, of which 9 were excluded from further analysis (6 samples were not suitable for the analysis and 3 samples were excluded after genetic identification as fox). Thus, we analysed 321 wolf scats (S4 Table), of which 100 were collected in the Poľana PLA (31%), 64 in the Vepor Mts (20%), 114 in the Muráň Plateau NP (36%), and 43 in the Poloniny NP (13%). The majority of the analysed scats, 94% (*n* = 302), contained remains of a single prey item, while 6% (*n* = 19) had two prey items. In total, we identified seven prey species in wolf scats: red deer, roe deer, wild boar and brown hare (*Lepus europaeus*) as wild prey, sheep as the only livestock species, and two unidentifiable species categorized as a rodent and a bird (Table 1).

**Table 1.** This is the Table 1 Diet composition. Winter diet composition of wolves in four study areas within the Slovak Carpathians during 2015 – 2017, measured as the percentage of occurrence (*FO*) and the percentage of the consumed biomass (*BM*). Is also shown the food resource category for each prey species in total as well as the n° of food items, n° of prey species and Niche Breath for each study area. ^a^ BM values < 0.1%.

| Study areas | Poľana PLA<br>(n=100) |  | Vepor Mts<br>(n=64) |  | Muráň Plateau NP<br>(n=114) |  | Poloniny NP<br>(n=43) |  | Total<br>(n=321) |  | Food resource<br>category [73] |
| --- | --- | --- | --- | --- | --- | --- | --- | --- | --- | --- | --- |
| Prey species | FO | BM | FO | BM | FO | BM | FO | BM | FO | BM |  |
| Red deer | 41.9 | 72.1 | 48.6 | 73.6 | 36.1 | 61.0 | 89.1 | 95.7 | 47.7 | 73.5 | Basic |
| Wild boar | 29.5 | 21.0 | 37.1 | 23.3 | 47.9 | 33.5 | 8.7 | 3.9 | 34.7 | 22.1 | Basic |
| Roe deer | 25.7 | 6.9 | 12.9 | 3.0 | 14.3 | 3.7 | 2.2 | 0.4 | 15.9 | 3.8 | Constant |
| Wild ungulates total | 97.1 | 99.9 | 98.6 | 99.9 | 98.3 | 98.2 | 100 | 100 | 98.2 | 99.4 | - |
| Sheep | 0.0 | 0.0 | 0.0 | 0.0 | 1.7 | 1.8 | 0.0 | 0.0 | 0.6 | 0.6 | Opportunistic |
| Brown hare | 0.0 | 0.0 | 1.4 | 0.1 | 0.0 | 0.0 | 0.0 | 0.0 | 0.3 | +<sup>a</sup> | Opportunistic |
| Rodent | 1.9 | +<sup>a</sup> | 0.0 | 0.0 | 0.0 | 0.0 | 0.0 | 0.0 | 0.6 | +<sup>a</sup> | Opportunistic |
| Bird | 1.0 | +<sup>a</sup> | 0.0 | 0.0 | 0.0 | 0.0 | 0.0 | 0.0 | 0.3 | +<sup>a</sup> | Opportunistic |
| Number of food items | 105 |  | 70 |  | 119 |  | 46 |  | 340 |  | - |
| Number of prey species | 5 |  | 4 |  | 4 |  | 3 |  | 7 |  | - |
| Niche Breath | 0.51 |  | 0.52 |  | 0.54 |  | 0.12 |  | 0.28 |  | - |

### Wolf diet composition

The majority of wolf diet was composed by wild ungulates (*FO* = 98.2%; *BM* = 99.4%). From the three most available wild ungulates within all study areas, red deer was the dominant prey (*FO* = 36.1% to 89.1%; *BM* = 61.0% to 95.7%), followed by wild boar (*FO* = 8.7% to 47.9%; *BM* = 3.9% to 33.5%), and roe deer (*FO* = 2.2% to 25.7%; *BM* = 0.4% to 6.9%)(Table 1). Given the *FO and* considering all study areas, red deer and wild boar are the only basic food resources for wolves in the Slovak Carpathians. Sheep was the only livestock species and was found in only two samples from the Muráň Plateau NP (*FO* = 1.7%; *BM* = 1.8%; Table 1). Small-sized prey, such as brown hare, rodents and birds had a marginal occurrence in wolf diet (*FO* < 2%) and accounting for less than 0.1% of the *BM* (Table 1).

Although the *FO* for red deer and wild boar showed no statistical differences (*χ*^2^ = 0.04; P = 0.851), the proportion of *BM* of red deer was significantly higher compared to wild boar (*χ*^2^ = 36.71; P < 0.001). Roe deer, categorized as a constant wolf prey (*FO* = 15.9%; *BM* = 3.8%), had significantly lower *FO* than the ones reported for red deer and wild boar (red deer: *χ*^2^ = 34.55; P < 0.001; wild boar *χ*^2^ = 0.04; P = 0.851), with a similar pattern for proportion of *BM* (red deer: *χ*^2^ = 140.48; P < 0.001= 0.851; wild boar: *χ*^2^ = 40.89; P < 0.001). There were considerable differences in the composition of prey between the four study areas. Red deer was the most frequent prey in Poloniny NP (*FO* = 89.1%), Poľana PLA (*FO* = 41.9%) and Vepor Mts (*FO* = 48.6%), however was significantly higher than wild boar only in the Poloniny NP (*χ*^2^ = 54.97; *P* < 0.001). In contrast, wild boar was the most frequent species in Muráň Plateau NP (*FO* = 47.9%), followed by red deer, although with no significant differences. In terms of *BM*, the pattern of red deer being dominant prey was consistent across all study areas (Table 1), with *BM* of red deer being significantly higher compared to wild boar in Poľana PLA (*χ*^2^ = 52.21; *P* < 0.001), Vepor Mts (*χ*^2^ = 33.16; *P* < 0.001), Muráň Plateau NP (*χ*^2^ = 17.22; *P* < 0.001) and Poloniny NP (*χ*^2^ = 71.63; *P* < 0.001). Roe deer was most consumed in the Poľana PLA (*FO* = 25.7%) while had much smaller occurrence in the Poloniny NP (*FO* = 2.2%).

The niche breadth was comparable for Poľana PLA, Vepor Mts and Muráň Plateau NP, with relatively high values ranging between 0.51 – 0.54, while wolves in the Poloniny NP had surprisingly narrow niche breadth (0.12) by feeding mostly on red deer, reflecting their dietary specialization (Table 1).

### Wolf prey selection

Compared to the available prey biomass, overall, wild boar occurred in wolf diet more frequently than expected (*D* = 0.31; Fig 3), while red deer use was proportional to its availability (*D* = −0.05; Fig 3), and roe deer was used less than available (*D* = −0.52; Fig 3). This trend was consistent across the Poľana PLA, Vepor Mts and Muráň Plateau NP, except for the Poloniny NP where wolves strongly selected for red deer (*D* = 0.81; Fig 3) and avoided wild boar (*D* = −0.62; Fig 3).

**Fig 3.**
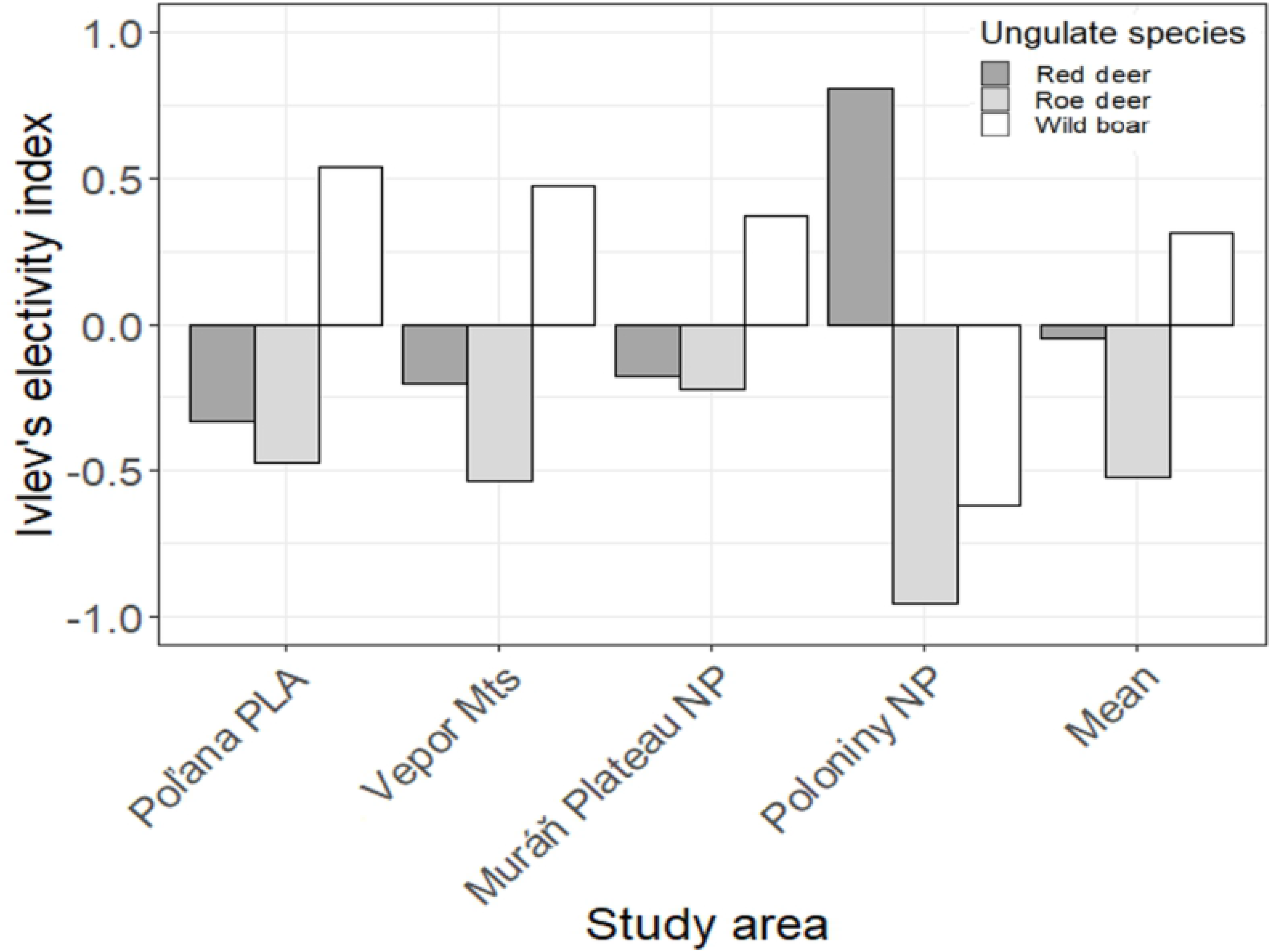
This is the Fig 3 Title: Ivlev’s index. This is the Fig 3 legend: Prey electivity (Ivlev’s index) for the main three wild ungulates according to the biomass consumed by wolves and the biomass available in each of the study areas in the Slovak Carpathians.

Wolves selected for red deer when red deer density increased (Fig 4a), and for wild boar when wild boar density increased (Table 2; Fig 4d), but this trend was stronger for the former prey species as the LSD coefficient for red deer (β = 1.95 ± 0.49; P < 0.001) was higher compared to wild boar (β = 1.02 ± 0.45; P = 0.02; Table 2). Roe deer was not selected based on its density (β = 0.66 ± 0.35; P = 0.06), but after applying density of other prey (red deer and wild boar; S3 Table), the probability of roe deer being found in a wolf scat decreased significantly with increasing density of red deer and wild boar (β = −2.79 ± 1.40; P = 0.05; Fig 4g). Elevation was a significant factor for all wild ungulates, however with differing direction and magnitude of selection (Table 2; Fig 4b, 4e and 4h). In particular, the probability for red deer being consumed by wolves increased at low elevations (β = −0.002 ± 0.001; P = 0.002), as well as for roe deer (β = −0.003 ± 0.001; P = 0.03), while the probability for wild boar consumption increased at high elevations (β = 0.003 ± 0.001; P < 0.001). Further, with increasing proportion of area with deep snow, the probability of being found in a wolf scat decreased for red deer (β = −0.25 ± 0.07; P < 0.001; Fig 4c), while it increased significantly for wild boar (β = 0.28 ± 0.08; P < 0.001; Fig 4f) and was not significant for roe deer (β = 0.001 ± 0.04; P < 0.74). Instead, we found significant differences in the probability of roe deer being found in a wolf scat between our study areas (Fig 4i; Table 2). Results of K-fold evaluations indicated that top models predicted wolf winter diet reasonably well (red deer ρ > 0.80; wild boar ρ > 0.92; roe deer ρ > 0.85).

**Fig 4.**
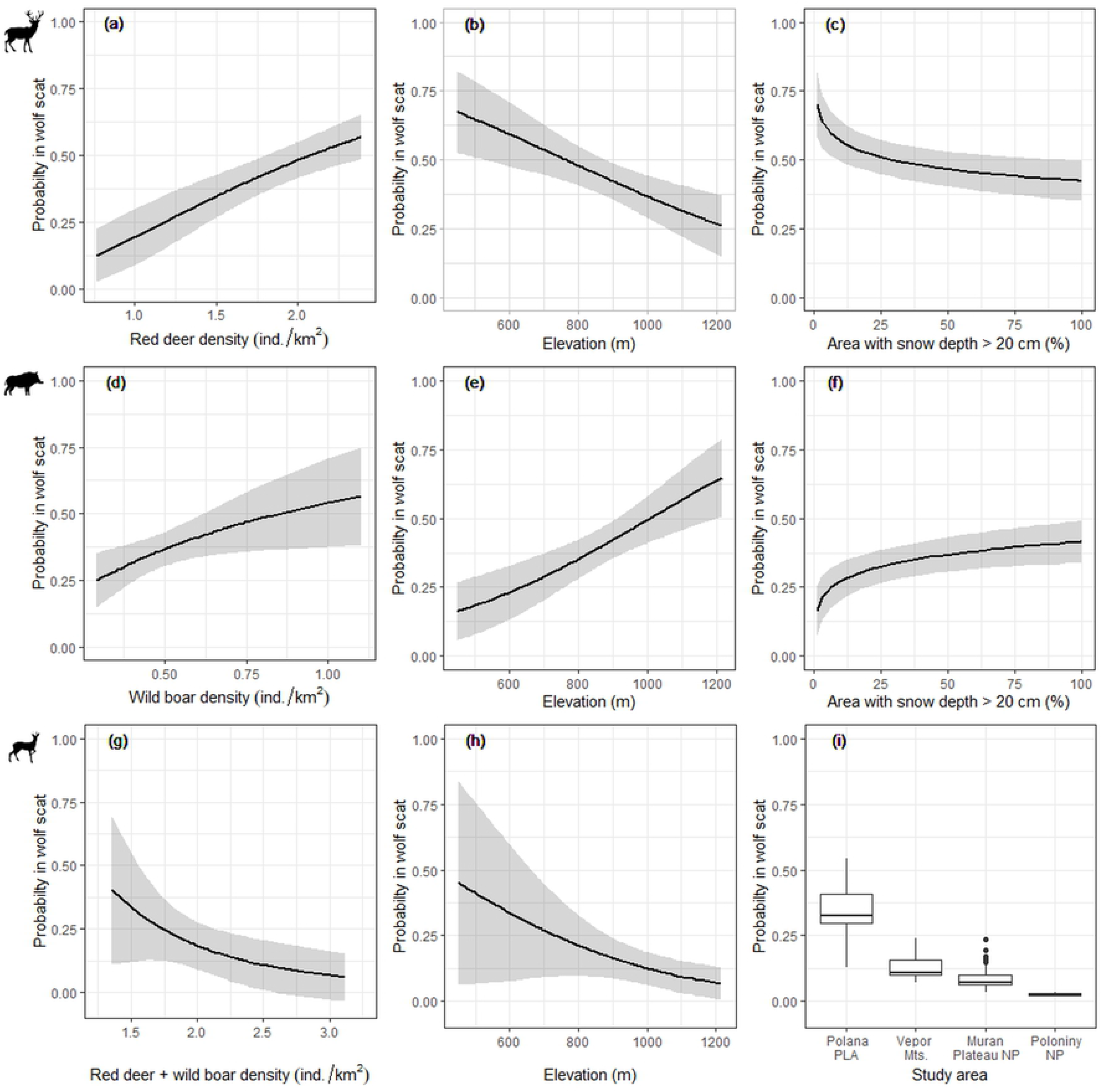
This is the Fig 4 Title: Latent selection difference. This is the Fig 4 legend: **Figure 4**. Predictions of the latent selection difference (LSD) for red deer (a, b, c), wild boar (d, e, f) and roe deer (g, h, i) being found in a winter wolf scat during 2015 – 2017 in the Slovak Carpathians, as a function of prey density, elevation, proportion of area with deep snow and a study area.

**Table 2.** This is the Table 2 Title: Latent selection difference. Latent selection difference (LSD) coefficients and confidence intervals of the top models for the main prey species as red deer, wild boar and roe deer, based on wolf scats collected during 2015 – 2017 in the Slovak Carpathians. Models describe the probability that one prey species would be selected in winter by wolves over another two prey species.

| Variable | Red deer |  |  | Wild boar |  |  | Roe deer |  |  |
| --- | --- | --- | --- | --- | --- | --- | --- | --- | --- |
| | $\beta$ | CI<sub>2.5%</sub> | CI<sub>97.5%</sub> | $\beta$ | CI<sub>2.5%</sub> | CI<sub>97.5%</sub> | $\beta$ | CI<sub>2.5%</sub> | CI<sub>97.5%</sub> |
| Intercept | 1.59 | 0.30 | 2.88 | -3.50 | -5.03 | -1.97 | - | - | - |
| log(Dens<sub>red deer</sub>) | 1.95 | 0.99 | 2.91 | - | - | - | - | - | - |
| log(Dens<sub>wild boar</sub>) | - | - | - | 1.02 | 0.14 | 1.90 | - | - | - |
| log(Dens<sub>red + boar</sub>) | - | - | - | - | - | - | -2.79 | -5.53 | -0.05 |
| Elevation | -0.002 | -0.003 | -0.001 | 0.003 | 0.001 | 0.005 | -0.003 | -0.006 | -6E-05 |
| log(Deep snow) | -0.25 | -0.40 | -0.10 | 0.28 | 0.12 | 0.44 | - | - | - |
| Location |  |  |  |  |  |  |  |  |  |
| Murán PNP | - | - | - | - | - | - | 3.18 | -0.23 | 6.59 |
| Poloniny NP | - | - | - | - | - | - | 0.19 | -0.72 | 1.10 |
| Poľana PLA | - | - | - | - | - | - | 4.58 | 3.21 | 5.95 |
| Vepor Mts | - | - | - | - | - | - | 3.31 | 2.17 | 4.45 |

## Discussion

The winter diet of Slovak wolves in the Carpathians is characterized by a high occurrence of wild large-sized and medium-sized ungulates, while other food categories, including livestock, comprised a negligible fraction of the diet. Among wild ungulates, red deer occurred most often in wolf scats, consistent with its highest density compared to other wild ungulates except in Poloniny NP where roe deer has the highest density. However, by comparing relative use to availability of wild prey, we found that wolves in fact selected for wild boar over red deer consistently in all study areas except for the Poloniny NP. We found that this inconsistency can be linked to changing prey vulnerability, which is context-dependent and has useful implications in the management of both wild ungulates and wolves.

### Wolf diet composition

Our study showed that wild ungulates composed 98.2% of wolf winter diet in Slovak Carpathians, over an insignificant occurrence of livestock and other wild species. In general, red deer was the basic wolf prey *sensu* Ruprecht [73] with the highest occurrence in wolf scat, followed by wild boar also a basic wolf prey, and by roe deer as a constant food resource. This is consistent with the diet of wolves living in natural habitats with abundance of wild prey [7], including in north-central Europe, as southern Sweden [9], northeast Germany [80] and throughout north-eastern to south-eastern Poland [10,19]; but also in the southern European regions, such as the western Italian Alps [81], northern Italy [82–84], northern and southern Spain [85,86] and northeast Portugal [13]. The similarity of these results together with our findings support the idea that in general wolves prefer wild prey over domestic species [8].

Wolf diet differed considerably in the composition of the wild ungulates between our study areas. In particular, red deer was the most prevalent prey in the Poľana PLA, in the Vepor Mts and especially in the Poloniny NP, while wild boar was the most consumed in the Muráň Plateau NP despite having one of the lowest wild boar densities among our study areas. Besides there where evidences of trophic specialization in a single prey species (red deer) as reflected by the narrow niche breadth in Poloniny NP. Our results also suggest the preference of local wolf populations to either wild boar (in Muráň Plateau NP) or red deer (in Poloniny NP), while roe deer was always avoided, being less consumed than its availability. These findings support the idea that wolf diet might vary depending on the environmental context, which contradicts previous works showing that wolves hunt the most abundant prey as in Poland [87], Romania [6,88], Italy [82,89], Spain [86] and Croatia [21]. However, wolf diet during summer might be different because in other parts of the Carpathians, high consumption of wild boar has been observed in winter but dropped considerably during summer [48,90]. This diet pattern can be related to a higher availability of more vulnerable prey species, such as livestock grazing in mountain pastures.

Although roe deer was the least frequent wolf prey among wild ungulates and is frequently hunted by wolves with a similar frequency across Europe [7], we found some variation between our study areas. While roe deer was an opportunistic prey in Poloniny NP, it was categorized as a basic prey in the Poľana PLA. However, in regions where large-sized wild prey is less available, the roe deer can even become the most important prey for wolves such as in northwest Spain [86], in the western Italian Alps [91] or in the northeast Germany [80,92]. We found no consumption of potential prey species occurring at low densities, such as bison and beaver in Poloniny NP and horse in the Muráň NP, although these species are reported as a regular wolf prey in other European regions, namely horses in Romania and Portugal [6,20], bison in Poland [19], and beaver in Latvia [93]. Low consumption of these species in our study areas may be related to higher availability of the three main wild ungulates (red deer, roe deer, wild boar).

We found only 2 wolf scats containing sheep remains. Livestock in Slovakia is usually brought to low elevations during winter and is kept in barns until spring, thus usually not being available as a wolf prey during our sampling period. While livestock consumption could be higher during summer when it is grazing at mountain pastures, a previous study from nearby areas in northern Slovakia showed only 4.8 % frequency of occurrence of livestock in the wolf diet during mountain grazing season [48]. This suggests that even in periods with high vulnerability, livestock is only an opportunistic prey for wolves in Slovak Carpathians given the high availability of wild ungulates.

### Wolf prey selection

On average, only wild boar was observed to be selected by Slovak wolves, while red deer and roe deer were generally avoided, with this pattern being consistent throughout the Poľana PLA, Vepor Mts and Muráň Plateau NP. Yet, in many European studies, red deer was the preferred prey of wolves [10,19,84,94,95] which is consistent to our findings from the Poloniny NP located in the Eastern Carpathians. Since wolves balance the difficulty of killing prey with the benefit obtained, red deer has been shown to be the optimal-size prey for typical central European packs of 4 – 6 wolves [61]. However, the positive selection for wild boar in the topographically variable Poľana PLA, Vepor Mts and Muráň Plateau NP can be explained by wild boar being the most vulnerable prey species in winter and at higher elevations regardless of their actual abundance, as also observed in Poland [54,87], Romania [6,88] and Italy [24,91,96]. First, wild boars live in large family groups where the percentage of young individuals is higher than in other ungulates and births are scattered over a longer period. Second, the distinct topography and higher altitudes of the Poľana PLA, Vepor Mts and Muráň Plateau NP with deeper snow may result in a considerable restriction of movement and depletion of foraging resources for wild boar [35,61], while red deer commonly winters and survives in such higher altitudes [69]. As a consequence, wolves are more likely to encounter vulnerable individuals of wild boar and potentially increase the relative predation success [24].

In contrast, the observed high selection for red deer in the Poloniny NP can be related to a low extent of highly productive open habitats such as pastures and agricultural land, which could make red deer more spatially predictable and vulnerable to wolf predation [81,91]. Simultaneously, the smallest elevation range and highest proportion of deciduous forests in Poloniny NP compared to the other study areas, can make wild boars a less vulnerable prey since they may benefit in term of forage availability, becoming with increased fitness and scatter throughout the entire area.

Given that availability of red deer and wild boar were similar between study areas, we conclude that a differential selection of wolves between central and eastern Slovakia might have been driven by distinct vulnerability of prey species induced by different environmental conditions rather than a display of the true prey switching (*sensu* Murdoch) [26] which is driven by changes in prey density. However, according to our predictions, Slovak wolves tend to increase consumption of red deer or wild boar when the population of each of these species increases, which shows their potential to switch the selection between these two ungulate species when the availability of one becomes higher than the other [25].

Roe deer on the other hand was constantly avoided across all our study areas in conformity to previous studies from Poland [10,87,95] and Italy [98], where this species occur within a diverse and abundant community of wild ungulates. Due to its small size and agility roe deer may not constitute a profitable prey and wolves should only prey on roe deer opportunistically when encountered [25]. Our results also confirmed this hypothesis because roe deer was more avoided in Poloniny NP despite this study area being the one where roe deer densities are higher than other larger and more profitable wild ungulates, such as red deer and wild boar.

Our results together with other examples across Europe illustrate extremely variable and flexible food habits of wolves across their range, with the capacity to exploit different vulnerability of their prey and also to swich their diet between areas following spatio-temporal variations in prey availability [24,25]. The observed context-dependent vulnerability of wild ungulates confirmed our hypothesis that there are indirect effects of environment on winter prey vulnerability through limitation of mobility and resources [25]. However, it would be interesting to develop further studies during summer in order to evaluate potential changes in prey selection by Slovak wolves and the influence of other environmental or ecological traits, such as water availability and livestock vulnerability.

### Management implications

Wolf predation is known to have an important sanitation effect on prey populations, by limiting the spread or incidence of fatal wildlife diseases, such as Anthrax in north American Bison (*Bison bison*) [100] or tuberculosis on Iberian wild boars [101]. In this context, the strong selection of wolves for wild boar in central Slovakia suggests that this carnivore may have the potential to eliminate individuals infected by the African swine fewer that is spreading throughout Slovakia, since 2019 [102]. This was strongly suggested during the latest epidemic of swine fever in Slovakia during 1994 – 2003, when the vast majority of the positive cases were identified outside of the wolf distribution range [103]. Furthermore, the presence of wolves can induce behavioural changes in wild ungulates through a landscape of fear [104], with the potential of reducing damages on agricultural crops [105] and young forest stands [104], which is highly relevant given the rapidly increasing numbers of wild ungulates throughout Slovakia and Europe. Finally, wolf predation on wild ungulates is often considered an economical damage and raises conflicts with hunters, although in Slovakia hunting grounds with losses on game species due to predators are annually compensated by the Ministry of Environment. However, considering our findings together with the available knowledge on wolf-prey interactions, the demographic impact of wolf predation on the abundant Slovak populations of wild ungulates may not be so significant and can have the potential to improve fitness of wild ungulates and reduce the risk of wolf damages to livestock. Consequently, given the frequent damages caused by wild ungulates in agriculture, forestry and road traffic [106], the positive role of large carnivores hunting their natural prey should be properly considered to achieve a future coexistence between wolves and human activities.

## Acknowledgements

We are thankful to all people that helped in field work for sample collection and in laboratory assistance for scat analysis. We thank the administrations of the Muráň Plateau NP, Poloniny NP, Poľana PLA and Lesy SR Hnúšťa for permits and logistic support related to field sampling.

## Supporting information

**S1_Table. This is the S1 Table Title.** This is the S1 Table legend: **S1 Table**. Estimated population sizes (number of individuals) of the main wild ungulate species within our study areas, Slovakia. (S1Table.pdf)

**S2_Table. This is the S2 Table Title.** This is the S2 Table legend: **S2 Table**. Average population sizes (number of individuals) and densities (individuals/km2) of the main livestock species during 2015 – 2017 in all municipalities located within our study areas, Slovakia. (S2Table.pdf)

**S3_Table. This is the S3 Table Title**. This is the S3 Table legend: **S3 Table**. Candidate logistic regression models explaining the probability that one species (red deer, wild boar, roe deer) would occur in winter wolf scat over other two species of wild ungulates. k – number of model components; LL – log-likelihood; AICc – Akaike Information Criterion for small sample sizes; w – AIC weight (best model highlighted in bold). (S3Table.pdf)

**S4_Table. This is the S4 Table Title**. This is the S4 Table legend: **S4 Table**. Data set information used in the analysis: sampling date (Month and Year) and presence of prey in each sample.

